# Evaluation of the microbial community structure of potable water samples from occupied and unoccupied buildings using16S rRNA amplicon sequencing

**DOI:** 10.1101/2020.07.17.209346

**Authors:** Kimothy L Smith, Howard A Shuman, Douglas Findeisen

## Abstract

We conducted two studies of water samples from buildings with normal occupancy and water usage compared to water from buildings that were unoccupied with little or no water usage due to the COVID-19 shutdown. Study 1 had 52 water samples obtained *ad hoc* from buildings in four metropolitan locations in different states in the US and a range of building types. Study 2 had 36 water samples obtained from two buildings in one metropolitan location with matched water sample types. One of the buildings had been continuously occupied, and the other substantially vacant for approximately 3 months. All water samples were analyzed using 16S rRNA amplicon sequencing with a MinION from Oxford Nanopore Technologies. More than 127 genera of bacteria were identified, including genera with members that are known to include more than 50 putative frank and opportunistic pathogens. While specific results varied among sample locations, 16S rRNA amplicon abundance and the diversity of bacteria were higher in water samples from unoccupied buildings than normally occupied buildings as was the abundance of sequenced amplicons of genera known to include pathogenic bacterial members. In both studies *Legionella* amplicon abundance was relatively small compared to the abundance of the other bacteria in the samples. Indeed, when present, the relative abundance of *Legionella* amplicons was lower in samples from unoccupied buildings. *Legionella* did not predominate in any of the water samples and were found, on average, in 9.6% of samples in Study 1 and 8.3% of samples in Study 2.

**Synopsis:** Comparison of microbial community composition in the plumbing of occupied and unoccupied buildings during the COVID-19 pandemic shutdown.

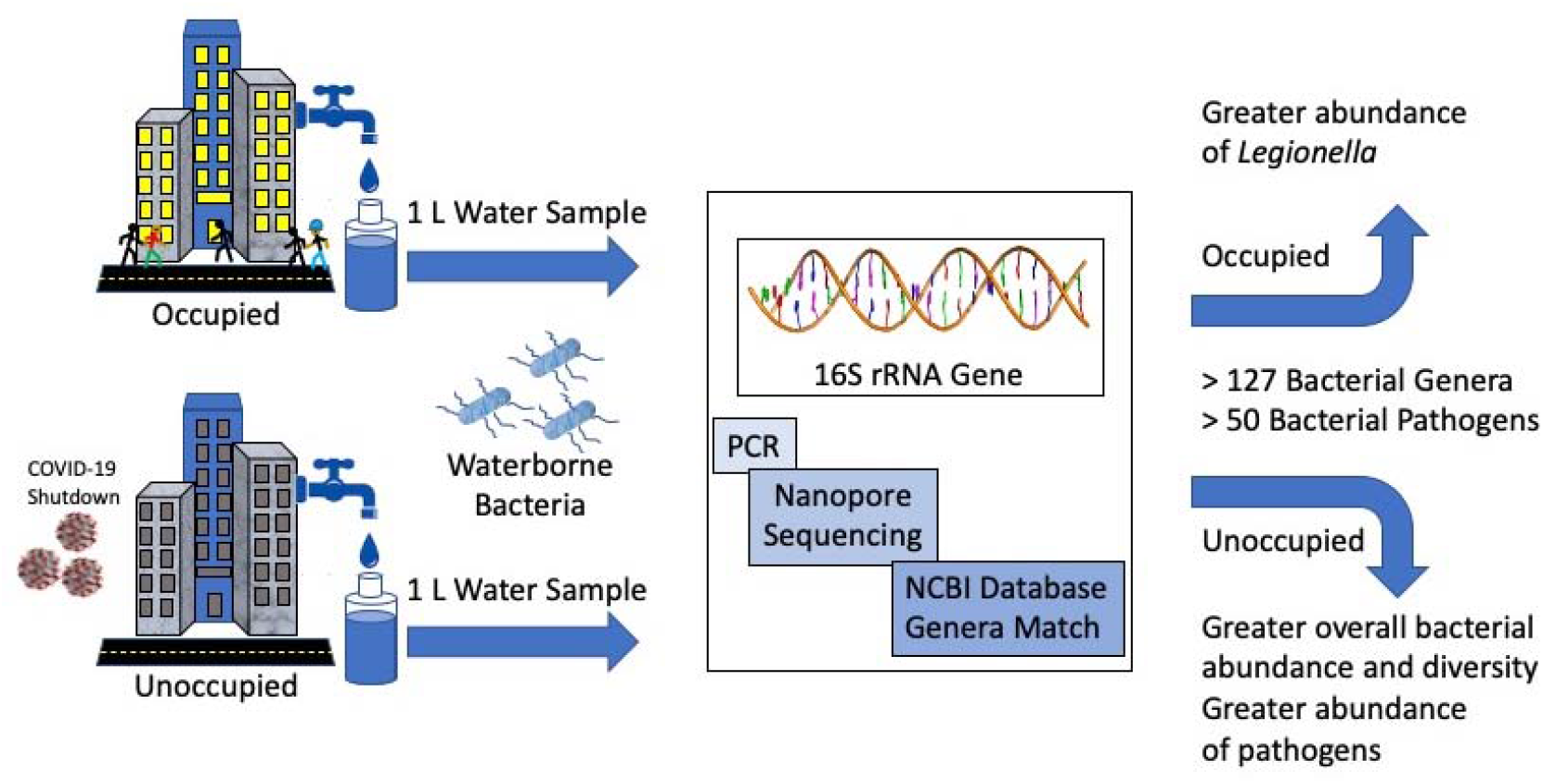

## Introduction

Stagnation of water in building plumbing systems can result in deterioration of water quality and proliferation of pathogens causing a major public health concern ^1^. Stagnation can also result in low or undetectable levels of residual disinfectant, such as chlorine ^2^. During the COVID-19 pandemic, the United States has experienced building shutdowns on a scale and duration not previously experienced. This disuse may present a risk to public health from pathogenic microbes that can grow in building plumbing systems ^2,3^. Numerous recently-issued guidance documents address the potential adverse effect on the quality of water that has stagnated in premise plumbing systems due to reduced occupancy during the current COVID-19 pandemic ^2,4–6^. Much of this guidance centers on *Legionella*, a plumbing-associated waterborne pathogen.

*Legionella* are found throughout the natural aquatic environment ^7^. *Legionella* growth in premise plumbing systems is supported by certain physical-chemical conditions that are associated with stagnant water: accumulated sediment, tepid temperatures, excessive water age and absence of residual disinfectant. Many of the recent guidance documents addressing re-occupancy of buildings after COVID-related closures regard *Legionella* as the predominant, if not the only pathogen of concern in stagnant water in under-occupied buildings; some assume that significant *Legionella* contamination is probable if not inevitable, and some suggest that testing only for *Legionella* is sufficient to determine whether or not a building is safe to re-occupy. Implementation of the recommendations in these guidance documents has important financial and public health implications. At the most fundamental level, we believe that the rationale for focusing on *Legionella* merits evidence-based scrutiny. One approach to this scrutiny is to characterize the microbial community structures of potable water from building plumbing systems, in order to identify bacterial genera with at least some members that are known to be pathogenic.

Marker gene amplicon sequencing is commonly used to assess microbial communities and has a nearly 40-year history ^8^. Molecular methods have distinct advantages over culturing and isolation; the routine isolation of all pathogens from water samples is impractical and cannot accurately characterize the microbial diversity in natural environments ^9,10^. Interestingly, despite public health relevance of potable water, very little information is available on metagenomic analyses of potable water systems ^11^. The 16S rRNA gene is a suitable choice for characterizing microbial community structures ^8^ and when combined with Next Generation Sequencing (NGS) methods, such as the Oxford Nanopore Technologies MinION™, can provide rapid and cost effective results.

The use of amplicon sequencing to assess microbial community composition and diversity is limited to identification at the genus level. In contrast to qPCR, amplicon sequencing does not broadly provide resolution to the species level. Rather, it provides a powerful means of initial screening for genera known to have at least some pathogenic members and can help elucidate the changes in and differences between microbial communities in potable water from different sources, such as occupied vs. unoccupied buildings. Studies like this one using 16S rRNA gene sequencing of potable water can provide useful and important insights that provide evidence-based observations to inform further, targeted testing necessary for the safe reopening of buildings after prolonged unoccupancy.

In this report we present the results of two studies (Study 1 and Study 2) of microbial communities in water from buildings with normal occupancy and water usage compared to water from buildings that were unoccupied with little or no water usage due to the COVID-19 shutdown. Study 1 had 52 water samples obtained from buildings in four metropolitan locations in different states in the US and a range of building types. Study 2 had 36 water samples obtained from two buildings in one metropolitan location with matched water sample types. All water samples were analyzed using 16S rRNA amplicon sequencing.

## Experimental Methods

For Studies 1 and 2 we collected a total of 88 1-liter water samples for analysis and comparison. Samples were taken from (a) buildings with low or no occupancy (“unoccupied”) for which flushing had not been implemented and (b) buildings with normal, uninterrupted occupancy (“occupied”).

For Study 1 we collected water samples from 52 buildings across 4 states— Michigan, Nevada, New York and Texas. Types of buildings sampled included residential condominiums, office buildings, hotels and dormitories. 1-L bulk water samples were collected from multiple locations in each building sampled. The number of samples and sample type—e.g., hot water or cold water, proximal or distal—varied from building to building.

For Study 2 we collected 36 water samples from two closely situated, very similar dormitory buildings on a college campus; one was occupied and the other unoccupied. Water sampling locations from the two buildings were similar; samples were taken from the main water supply trunks, hot water supply, and from locations distributed spatially throughout the buildings. The number, type and location of samples were matched as closely as possible.

Samples were shipped overnight in coolers to help maintain stable temperature. Sample processing, initiated within 24 hours of receipt, was as follows:

- <u>Concentration</u> Samples were concentrated through a membrane ultrafilter (*Filpath*™ Ultrafilter, Nephros, Inc., South Orange, NJ, USA)
- <u>Lysis</u> Sample concentrate remaining on the filter was treated with a lysis buffer solution (Environmental Lysis Solution, Chai, Inc., Santa Clara, CA) and then removed.
- <u>Amplification</u> Lysate samples were added directly to PCR reaction mixes for amplifying the 16S rRNA genes. Amplification of 16S rRNA genes was conducted using the 16S Barcoding Kit (SQK-RAB204; Oxford Nanopore Technologies, Oxford, UK) containing the 27F/1492R primer set, LongAmp™ Taq 2x Master Mix (New England Biolabs, Ipswich, MA, USA), and Chai Green Dye 20x (Chai, Inc., Santa Clara, CA, USA). Amplification was performed using a Chai Open™ qPCR thermal cycler (Chai, Inc., Santa Clara, CA, USA) with the following PCR conditions: initial denaturation at 95 °C for 30 s, 40 cycles of 95°C for 30 s, 53°C for 30 s, and 65°C for 2 min.
- <u>Library preparation</u> PCR products were purified using AMPure XP (Beckman Coulter, Indianapolis, IN, USA), two 70% alcohol washes, and quantified by a NanoDrop (Thermo Fischer Scientific, USA). Purified amplicons were pooled (up to 12) and a total of 100 ng DNA was used for library preparation.
- <u>Amplicon sequencing</u> MinION™ sequencing was performed using R9.4 flow cells (FLO-MIN106; Oxford Nanopore Technologies) according to the manufacturer’s instructions. MINKNOW software ver. 19.12.5 (Oxford Nanopore Technologies) was used for data acquisition.
- <u>Bioinformatic analysis</u> The EPI2ME™ platform 16S Workflow (Metrichor, Ltd., a subsidiary of Oxford Nanopore Technology) was used to classify sequencing reads and perform bacterial genera identification.
- <u>Estimates of bacterial concentrations</u> Estimates of bacterial concentrations (as CFU/mL) were based on Cq values for each 1-liter water sample by (a) estimating the starting 16S rRNA target DNA copies present in the PCR reaction volume, and then performing a series of back-calculations estimating the number of CFU per milliliter of bacteria present in each of the 1-liter samples of water concentrated on the filters, assuming one copy of the template target per cell.

## Results and Discussion

Table 1 and 2 provide water sample information collected for Study 1 and 2, respectively. The measured DNA concentration (Study 1 only) was measured using the lysate (post-filter concentration and lysis) using a Thermo-Fisher NanoDrop™ 1000. This DNA concentration is all-source (prokaryotic and eukaryotic) in contrast to the 16S rRNA gene sequence abundance which is representative of bacterial DNA presence in the samples.

**Table 1.** Study 1 Sample Information

| Sample # | Unoccupied/<br>Occupied | State | Date<br>Collected | Location | Description | Temp | Chlorine<br>ppm | DNA<br>ng/ul | 16S rRNA<br>Gene<br>Amplicon<br>Sequence<br>Abundance<br>(Reads) |
| --- | --- | --- | --- | --- | --- | --- | --- | --- | --- |
| 1 | Occupied | NY | 5/6/20 | Bath shower - hot | First draw | 110 deg F |  | 6.9 | 4 |
| 2 | Occupied | NY | 5/6/20 | Tub | First draw | 110 deg F |  | 24.5 | 140 |
| 3 | Occupied | NY | 5/6/20 | Bath sink - cold | First draw | <60 deg F |  | 5.3 | 7 |
| 4 | Occupied | NY | 5/6/20 | Kitchen sink - Hot | First draw | 115 deg F |  | 6.4 | 17 |
| 5 | Unoccupied | NV | 5/7/20 | Top floor tower |  | Room<br>temp |  | 8.0 | 97,584 |
| 6 | Unoccupied | NV | 5/7/20 | Ground floor sink | First out | Room<br>temp |  | 6.8 | 54,076 |
| 7 | Unoccupied | NV | 5/7/20 | Ground floor sink |  | Room<br>temp |  | 8.6 | 52,260 |
| 8 | Unoccupied | NV | 5/7/20 | Top floor 7th |  | Room<br>temp |  | 13.2 | 154,259 |
| 9 | Occupied | NV | 5/12/20 | Mens room | 5 story building | Room temp |  | 4.7 | 63,013 |
| 10 | Occupied | NV | 5/12/20 |  |  | Room<br>temp |  | 3.9 | 14 |
| 11 | Occupied | NV | 2/12/20 |  |  | Room<br>temp |  | 7.9 | 92,421 |
| 12 | Unoccupied | NY | 5/13/20 | Kitchen sink | 36th floor |  |  | 27.7 | 671 |
| 13 | Unoccupied | NY | 5/13/20 | Kitchen sink | 45th floor |  |  | 14.2 | 34 |
| 14 | Unoccupied | NY | 5/13/20 | Kitchen sink | 36th floor |  |  | 26.6 | 11,450 |
| 15 | Unoccupied | NY | 5/13/20 | Kitchen sink | 45th floor |  |  | 16.1 | 108,430 |
| 16 | Unoccupied | NY | 5/12/20 | Bath sink - cold | Unit 4A | 60 deg F |  | 11.9 | 12 |
| 17 | Unoccupied | NY | 5/12/20 | Kitchen sink - Hot | Unit 4E | 70 deg F |  | 13.9 | 6 |
| 18 | Unoccupied | NY | 5/12/20 | Shower/tub - hot | Unit 9J | 95 deg F |  | 14.2 | 26 |
| 19 | Unoccupied | NY | 5/12/20 | Porter's closet -<br>cold | 10th floor | 65 deg F |  | 6.4 | 250,316 |
| 20 | Unoccupied | NY | 5/12/20 | Bath sink - hot | 4th floor | 70 deg F |  | 84.4 | 20 |
| 21 | Unoccupied | MI | 5/11/20 | Drinking fountain | 24th floor |  |  | 6.5 | 8,686 |
| 22 | Unoccupied | MI | 5/11/20 | Drinking fountain | Basement |  |  | 3.3 | 447 |
| 23 | Unoccupied | MI | 5/11/20 | Drinking fountain | 24th floor |  |  | 5.6 | 17,151 |
| 24 | Occupied | MI | 5/8/20 | Kitchen sink | 1st floor | 57 deg F |  | 2.3 | 1,274 |
| 25 | Occupied | MI | 5/8/20 | Kitchen sink | 1st floor | 57 deg F |  | 4.4 | 558 |
| 26 | Unoccupied | MI | 5/11/20 | Drinking fountain | Basement |  |  | 4.4 | 32,568 |
| 27 | Occupied | NY | 5/6/20 | Kitchen sink | First draw | 70 deg F |  | 7.5 | 3 |
| 28 | Occupied | NY | 5/6/20 | Bath sink - hot | First draw | 120 deg F |  | 13.3 | 5 |
| 29 | Unoccupied | TX | 5/15/20 | Manual sink - hot<br>Distilled | First floor | 110 deg F | 0.52 | 3.2 | 1,763 |
| 30 | Occupied | TX | 5/15/20 | Manual faucet -<br>cold |  | 76 deg F | 0.22 | 5.3 | 220,957 |
| 31 | Unoccupied | TX | 5/15/20 | Shower - cold |  | 79 deg F | 0.20 | 3.7 | 619 |
| 32 | Unoccupied | TX | 5/15/20 |  | First floor | 74 deg F | 1.22 | 5.0 | 1,360 |
| 33 | Occupied | TX | 5/15/20 | Manual faucet sink | 2nd floor | 77 deg F | 0.29 | 3.5 | 64,188 |
| 34 | Occupied | TX | 5/15/20 | Manual faucet | First floor | 75 deg F | 0.26 | 3.1 | 122 |
| 35 | Unoccupied | TX | 5/15/20 | Manual sink - hot proximal | 6th floor | 112 deg F | 0.72 | 2.7 | 43 |
| 36 | Unoccupied | TX | 5/15/20 | Sink manual | 5th floor | 73 deg F | 1.12 | 4.6 | 181 |
| 37 | Unoccupied | TX | 5/15/20 | Sink manual - hot proximal | 6th floor | 111 deg F | 0.78 | 2.5 | 33 |
| 38 | Unoccupied | TX | 5/15/20 | Sink - cold | 6th floor | 73 deg F | 1.21 | 4.1 | 64 |
| 39 | Occupied | TX | 5/15/20 | sink proximal - hot |  | 114 deg F | 0.14 | 2.4 | 177,696 |
| 40 | Occupied | TX | 5/15/20 | Sink - cold | 2nd floor<br>NW corner | 77 deg F | 0.28 | 3.0 | 844 |
| 41 | Unoccupied | TX | 5/15/20 | Manual sink - cold ambient | 5th floor | 74 deg F | 1.12 | 4.6 | 4 |
| 42 | Unoccupied | TX | 5/15/20 | Manual sink | First floor | 76 deg F | 1.48 | 3.4 | 995 |
| 43 | Unoccupied | TX | 5/15/20 | Manual sink - hot Distilled | First floor | 112 deg F | 0.68 | 3.1 | 11,604 |
| 44 | Occupied | TX | 5/15/20 | Dynamic flow sink - cold |  | 76 deg F | 0.24 | 3.3 | 586 |
| 45 | Unoccupied | TX | 5/15/20 | Manual sink - cold | 6th floor | 72 deg F | 1.32 | 4.1 | 304,833 |
| 46 | Occupied | TX | 5/15/20 | cold | First floor | 76 deg F | 0.27 | 3.1 | 20,092 |
| 47 | Occupied | TX | 5/15/20 | Sink Auto Faucet - hot | First floor | 114 deg F | 0.13 | 2.7 | 5,513 |
| 48 | Occupied | TX | 5/15/20 | cold | First floor | 78 deg F | 0.25 | 17.2 | 27 |
| 49 | Unoccupied | TX | 5/15/20 | Manual sink - cold dynamic | First floor | 74 deg F | 1.29 | 5.5 | 4,446 |
| 50 | Occupied | TX | 5/15/20 |  |  | 138 deg F | 0.14 | 4.6 | 2,538 |
| 51 | Unoccupied | TX | 5/15/20 | Dynamic | First floor | 70 deg F | 0.20 | 4.9 | 1,526 |
| 52 | Occupied | TX | 5/15/20 | Distilled |  | 138 deg F | 0.13 | 3.6 | 7 |

**Table 2.** Study 2 Sample Information

| **Sample #** | **Unoccupied/**<br>**Occupied** | **State** | **Date** | **Location** | **Description** | **Temp** | **Amplicon**<br>**Abundance** |
| --- | --- | --- | --- | --- | --- | --- | --- |
| 1 | Unoccupied | NY | 5/27/20 | Boiler Rm | CW@main - hose bib outside | 57 deg F | 26 |
| 2 | Unoccupied | NY | 5/27/20 |  | HW supply from tank - first draw | 124 deg F | 62 |
| 3 | Unoccupied | NY | 5/27/20 |  | Cold makeup for boiler | 76 deg F | 118,250 |
| 4 | Unoccupied | NY | 5/27/20 |  | HW supply flushed draw | 124 deg F | 203 |
| 5 | Unoccupied | NY | 5/27/20 | Suite 1116-1117 | Shower head hotside - first draw | 102 deg F | 40,613 |
| 6 | Unoccupied | NY | 5/27/20 | Suite 1116-1117 | Shower head hotside - flushed draw | 124 deg F | 11 |
| 7 | Unoccupied | NY | 5/27/20 | Suite 1140-1141 | Shower head - flushed draw | 122 deg F | 218 |
| 8 | Unoccupied | NY | 5/27/20 | Suite 1133-1134 | Shower head - first draw | 122 deg F | 170 |
| 9 | Unoccupied | NY | 5/27/20 | Suite 705-706 | Shower head hotside - first draw | 117 deg F | 28 |
| 10 | Unoccupied | NY | 5/27/20 | Room 735 | Shower head hotside - first draw | 122 deg F | 59 |
| 11 | Unoccupied | NY | 5/27/20 | Room 735 | Shower head hotside - flushed draw | 122 deg F | 188 |
| 12 | Unoccupied | NY | 5/27/20 | Suite 722-723 | Shower head hotside - first draw | 70 deg F | 711 |
| 13 | Unoccupied | NY | 5/27/20 | Suite 311-312 | Shower head cold - first draw | 73 deg F | 0 |
| 14 | Unoccupied | NY | 5/27/20 | Suite 311-312 | Shower head hotside - first draw | 123 deg F | 2 |
| 15 | Unoccupied | NY | 5/27/20 | Suite 338-339 | Shower head hotside - first draw | 115 deg F | 0 |
| 16 | Unoccupied | NY | 5/27/20 | Suite 318-319 | Shower head hotside - first draw | 75 deg F | 0 |
| 17 | Unoccupied | NY | 5/27/20 | Suite 318-319 | Shower head hotside - flushed draw | 121 deg F | 1 |
| 18 | Unoccupied | NY | 5/27/20 | Suite 1140-1141 | Shower head - first draw | 121 deg F | 58 |
| 19 | Occupied | NY | 5/27/20 | City Main | First draw | 57 deg F | 2 |
| 20 | Occupied | NY | 5/27/20 | City Main | 1 minute flush | 57 deg F | 4 |
| 21 | Occupied | NY | 5/27/20 | Boiler Rm | HW return top of tank - 10 second flush | 130 deg F | 374 |
| 22 | Occupied | NY | 5/27/20 | Boiler Rm | HW supply Heater #2 - First draw | 122 deg F | 1,123 |
| 23 | Occupied | NY | 5/27/20 | 4K? | Shower head hotside - first draw | 125 deg F | 1 |
| 24 | Occupied | NY | 5/27/20 | Apt 6F | Shower head hotside - first draw | 125 deg F | 1 |
| 25 | Occupied | NY | 5/27/20 | Apt 6F | Shower head hotside - flushed draw | 130 deg F | 571 |
| 26 | Occupied | NY | 5/27/20 | Apt 6O | Shower head hotside - first draw |  | 23 |
| 27 | Occupied | NY | 5/27/20 | Apt 6O | Shower head hotside - flushed draw | 138 deg F | 779 |
| 28 | Occupied | NY | 5/27/20 | Apt 6J | Shower head hotside - first draw | 123 deg F | 32 |
| 29 | Occupied | NY | 5/27/20 | Apt 4D | Shower head hotside - flushed draw | 70 deg F | 114 |
| 30 | Occupied | NY | 5/27/20 | Unit 4B | Shower head hotside - first draw | 98 deg F | 2,195 |
| 31 | Occupied | NY | 5/27/20 | 4B | Shower head hotside - flushed draw | 117 deg F | 5 |
| 32 | Occupied | NY | 5/27/20 | Unit 2F | Kitchen sink hot side - first draw | 73 deg F | 153,532 |
| 33 | Occupied | NY | 5/27/20 | Unit 1E | Shower head hotside - first draw | 78 deg F | 3,166 |
| 34 | Occupied | NY | 5/27/20 | Unit 1G | Shower head hotside - first draw | 140 deg F | 10 |
| 35 | Occupied | NY | 5/27/20 | Unit 1H | Shower head - first draw | 126 deg F | 15 |
| 36 | Occupied | NY | 5/27/20 | Unit 1H | Shower head - flushed draw | 137 deg F | 54 |

### Bacterial proliferation in stagnant water of unoccupied buildings

In Study 1 we generated a total of 1,765,493 16S rRNA gene amplicon sequences from the 52 water samples. The average amplicon abundance of unoccupied building samples was more than twice that of samples from occupied buildings, with the ratio of 16S rRNA gene amplicon abundance (unoccupied:occupied) ranging 1.5:1 to more than 1400:1. The microbial communities, as indicated by bacterial genera, in unoccupied building samples appear to be more diverse—i.e., 82% more bacterial genera are present. Both unoccupied and occupied samples contain multiple bacterial genera; 10-80% of the identified bacteria genera are putative potential human pathogens, based on the Centers for Disease Control and Prevention (CDC) list of Opportunistic Pathogens of Premise Plumbing ^12^. The amplicon sequence abundance of pathogenic bacteria appears to be generally higher in unoccupied samples, but the percentage proportion of amplicon sequence of pathogenic bacteria appears to be generally lower. In Study 2 we generated a total of 322,601 16S rRNA gene amplicon sequences form the 36 water samples. The average amplicon abundance of unoccupied building samples was almost 1.5 times that of occupied samples, with the ratio of 16S rRNA gene amplicon abundance (unoccupied:occupied) 6:1. Overall the combined 88 samples yield a ratio of amplicon abundance of 1.6:1 and an average amplicon abundance ratio of 1.4:1 (see Table 3).

**Table 3:** Ratios of Total and Average Amplicon Sequences for All Water Samples Unoccpuied:Occupied

|  | Unoccupied | Occupied | Ratio of Unoccupied:Occupied |
| --- | --- | --- | --- |
| Total Abundance of 16S rRNA gene Amplicon Sequences for All Samples | 1,275,448 | 812,646 | 1.57:1 |
| Average Abundance of 16S rRNA gene Amplicon Sequences for All Samples | 27,137 | 19,821 | 1.37:1 |

In both studies we used an intercalating dye in the PCR reaction when amplifying the 16S rRNA gene target and were only able to obtain a Cq value for 14 of the 52 samples in Study 1and 7 of the 36 samples in Study 2. For the 14 samples in Study 2 with a Cq value we calculated an estimate of the total bacterial bioburden, or the CFU/ml inclusive of all bacteria in the samples. The calculated CFU/ml values ranged from approximately 8 to approximately 2,000 CFU/ml. The average CFU/ml estimate for unoccupied building water samples in Study 1 was approximately 6 times higher than the average CFU/ml for occupied samples. From the 7 Cq values obtained in Study 2 the calculated CFU/ml values ranged from approximately 8 to approximately 1,280 CFU/ml. The number of occupied vs unoccupied samples with Cq values was not sufficient for a comparison. For those samples that did not yield a Cq value, we can only conclude that the bioburden was less than 2 CFU/ml in each of those samples.

### Bacterial pathogen proliferation in stagnant water of unoccupied buildings

For simplicity, in both Studies 1 and 2, only the 12 most abundant bacterial genera in each sample were considered for further analysis and comparison. The cutoff of 12 was arbitrarily chosen.

The total abundance of 16S rRNA gene amplicon sequences of in Study 2 for the 12 most abundant bacterial genera was 7,411. Table 4 shows the comparison of gene amplicon sequence proportions for pathogenic genera and nonpathogenic genera in occupied and unoccupied building samples.

**Table 4:**
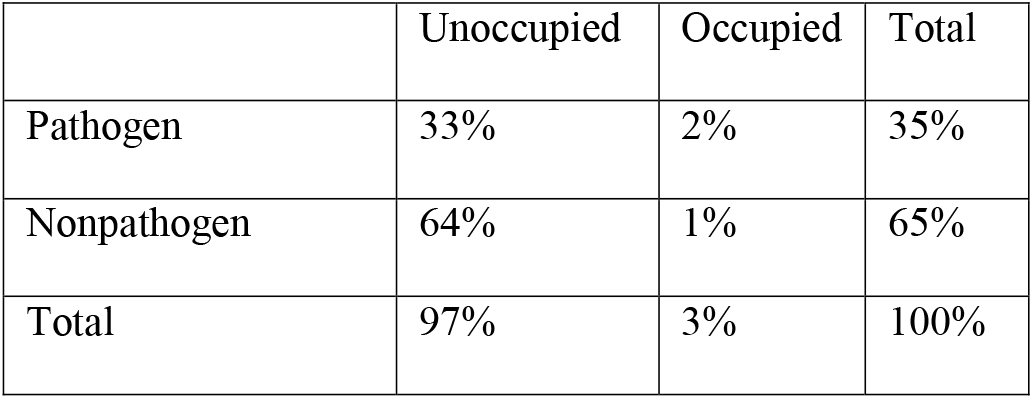
Proportions of 16S rRNA gene amplicon sequences among pathogen and nonpathogen genera in occupied and unoccupied building samples

The shift observed here of relative dominate proportion from occupied to unoccupied between pathogen and nonpathogen is similarly seem in Study 1. While pathogens seem to predominate in proportion in occupied building water samples when compared to nonpathogen genera, this is inverted in unoccupied building water samples. This phenomenon would seem to indicate a complex microbial community dynamic that merits further examination. Additionally, the proportions of specific bacterial genera gene amplicon sequences show changes or shifts between occupied and unoccupied building water samples (see Figure 1) also indicative of a dynamic microbial community composition.

**Figure 1:**
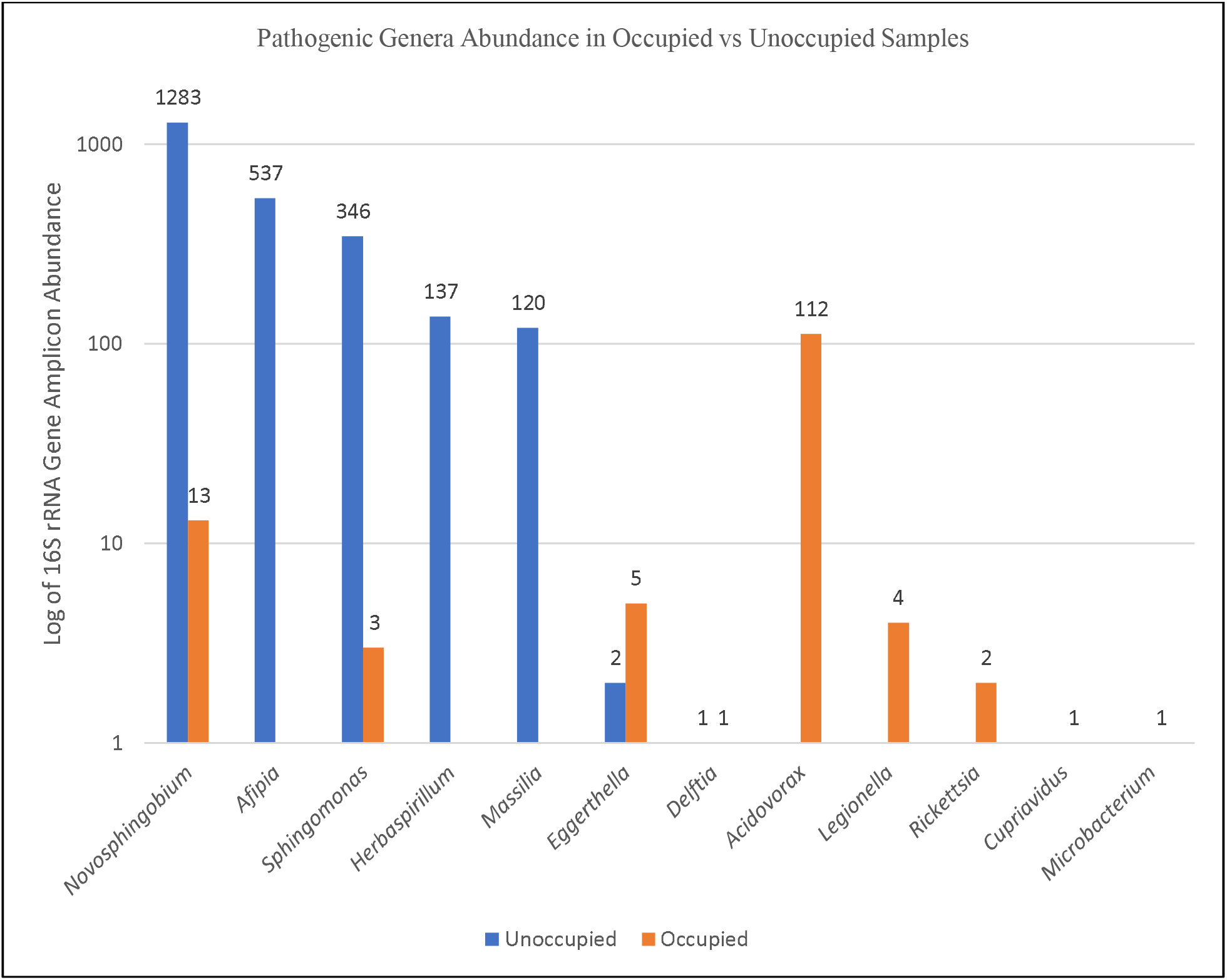
Log of putative pathogenic genera abundance in occupied vs unoccupied building water samples in Study 2 indicating a shift in microbial community composition.

### Presence and abundance of *Legionella* in the study samples

*Legionella sp.* was detected in 9.6% of all samples, including in 16.1% of unoccupied building water samples and 4.7% of non-stagnant samples. In samples where *Legionella sp.* was detected, it was less abundant than 59% of other pathogenic bacteria genera detected.

Study 1 had 5 samples that contained *Legionella*. Figure 2 shows the 16S rRNA gene amplicon percent abundance for the putative pathogen genera in those water samples.

**Figure 2:**
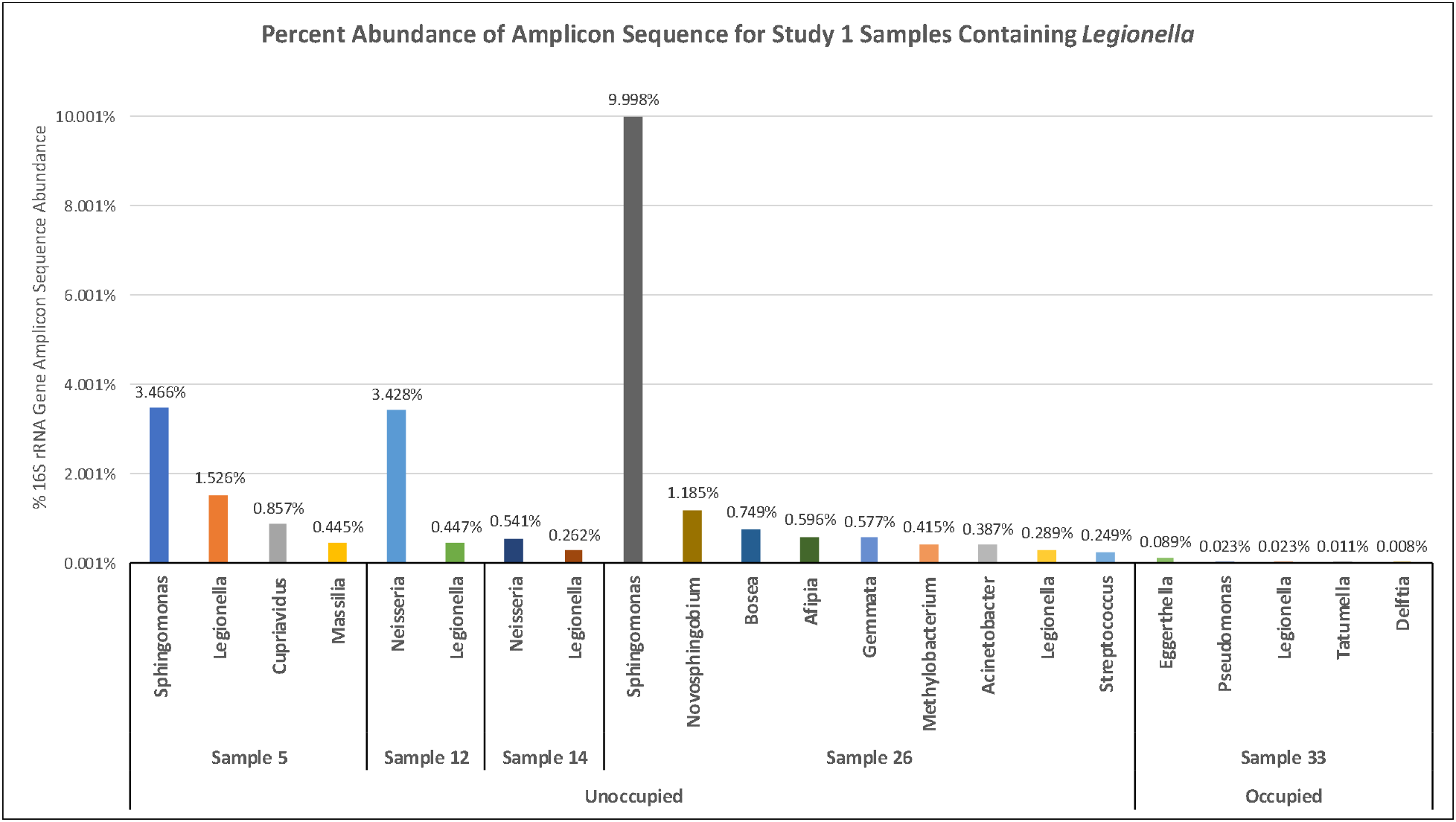
Percent abundance of 16S rRNA gene amplicon sequence for 5 samples from Study 1 (4 unoccupied water samples; 1 occupied water sample) which contained Legionella.

In Study 2, 3 samples contained *Legionella* at an exceedingly low level except one. A single occupied building water sample in Study 2 was taken from a currently unoccupied single living space that had a slow, constant leaking fixture (a kitchen sink faucet). This sample was removed from the further combined analyses for reasons given below but, presents a special case and deserves additional consideration. 16S rRNA gene amplicon sequencing of this sample resulted in the largest total number of reads for a single sample in the study (153,532; almost 50% of the grand total study-wide), and the second largest number of reads in the 12 most abundant genera selected for further analysis (81,513). This single sample accounted for almost 80,000 total putative pathogen sequenced amplicon, 320 of which were identified as *Legionella* (the largest abundance of *Legionella* of any sample in the Study by a factor of more than 100), and 2,333 nonpathogen reads.

## Conclusions

The results of these studies should be considered in light of the caveats mentioned previously and with consideration of the limitations described below. Notwithstanding the caveats and limitations, we believe the results have immediate practical implications with respect to the planning for re-occupancy of buildings following low occupancy due to COVID-19 measures (where water has stagnated in the pipes). Conventional wisdom and the scientific literature indicate that when potable water is allowed to dwell in building plumbing for a long period of time that stagnation will occur, and microorganisms will proliferate. It is likely that buildings that have been vacant or experienced low occupancy due to the recent or ongoing shutdowns will have greater levels of bacteria (including potential human pathogens) than expected under normal occupancy. Given the results presented in these studies water safety managers should not assume the water is contaminated with *Legionella* but rather that it likely harbors other pathogenic bacteria which present a public health hazard if exposed to or consumed by humans. Water testing only for *Legionella* is therefore insufficient and may provide false security against other significant health threats. If water samples are tested, they should be screened for multiple genera using technology similar to those described in these studies or equivalent, then tested for specific pathogens of concern based as indicated by the initial screen.

The results of these studies also underscore the complexity of microbial community dynamics that are resident in building premise plumbing systems. The sample taken in Study 2 from the leaky kitchen faucet also remind us that local effects – a constant, slow flow of nutrient with insufficient force to dislodge biofilms – can drastically impact the microbial community and ecosystem locally and lead to a potentially disastrous situation for someone consuming the water and variety of microbe. There are many questions yet to be addressed and answered utilizing evidence-based observations aided by current technologies.

### Study Limitations

∘ The buildings in Study 1 were not graded by degree of water usage or length of reduced occupancy; in some cases, these details are not known. Assigned categories were binary—unoccupied *vs.* occupied—and did not always consider potentially significant differences in building type, usage patterns, quality/variability in the water supply to the building, or other factors that could have a profound effect on test results.
∘ Sample analysis was not performed in triplicate.
∘ The precision of estimates of bacterial concentration (as CFU/mL) based on qPCR is inherently limited. Comparisons made of estimated bacterial populations in stagnant *vs.* non-stagnant samples should be considered qualitative. Estimates of bacterial concentrations were made for only 14 of the 52 samples.
∘ Classification of the microbial community structure based on comparing targeted sequencing data to databases of 16S rRNA can yield bacterial identity to the level of genera—i.e, operation taxonomic units (OTU) or amplicon sequence variants (ASV)—but may not be sufficiently reliable in all cases for identification at the species level.
∘ Comparisons between marker gene amplified sequence reads can be biased and confounded, particularly when attempting to draw conclusions regarding environmental or ecological effects.

## Acknowledgements

We wish to thank all of the participants in these study that provided water samples for analysis despite the risk of exposure to COVID-19.

